# Regularized-Ncut: Robust and homogeneous functional parcellation of neonate and adult brain networks

**DOI:** 10.1101/737627

**Authors:** Qinmu Peng, Minhui Ouyang, Jiaojian Wang, Qinlin Yu, Chenying Zhao, Michelle Slinger, Hongming Li, Yong Fan, Bo Hong, Hao Huang

**Author notes:** Corresponding Author: Hao Huang, Ph.D., 3401 Civic Center Blvd, Philadelphia, PA 19104.

## Abstract

Brain network parcellation based on resting-state functional MRI (rs-fMRI) is affected by noise, resulting in spurious small patches and decreased functional homogeneity within each network. Robust and homogeneous parcellation of neonate brain is more difficult with neonate rs-fMRI associated with higher level of noise and no functional atlas as spatial constraints. To meet these challenges, we developed a novel data-driven Regularized Normalized-cut (RNcut) method. RNcut is formulated by adding two regularization terms, a smoothing term using Markov random fields and a small-patch removal term, to conventional normalized-cut (Ncut) method. The RNcut and competing methods were tested with simulated datasets with known ground truth and then applied to both adult and neonate rs-fMRI datasets. Based on the parcellated networks generated by RNcut, intra-network connectivity was quantified. The test results from simulated datasets demonstrated that the RNcut method is more robust (p<0.01) to noise and can delineate parcellated functional networks with significantly better (p<0.01) spatial contiguity and significantly higher (p<0.01) functional homogeneity than competing methods. Application of RNcut to neonate and adult rs-fMRI dataset revealed distinctive functional brain organization of neonate brains from that of adult brains. Collectively, we developed a novel data-driven RNcut method by integrating conventional Ncut with two regularization terms, generating robust and homogeneous functional parcellation without imposing spatial constraints. A broad range of brain network applications and analyses, especially exploratory investigations of parcellating neonate and infant brain with noisy dataset, can potentially benefit from this RNcut method.

## Introduction

It has been reproducibly found that spontaneous low-frequency fluctuations in brain activities, monitored by blood oxygen level-dependent (BOLD) signals using resting-state functional magnetic resonance imaging (rs-fMRI), were temporally correlated across functionally related areas (Friston 1994; Biswal et al., 1995). Functional connectivity based on temporal correlation of BOLD signals is critical for characterizing intrinsic brain functional organization, which can be represented by parcellated networks (Bullmore and Sporns, 2009). Each brain network is located in functionally specific brain region. Based on these parcellated functional networks (FNs), distinguished organizational principles in the normal (e.g. Smith et al., 2009) and diseased brain (e.g. Bassett et al., 2008; Yerys et al., 2017) can be identified. Understanding individual variability of parcellated cortical regions in normal brains can facilitate delineating disease susceptibility (Mueller et al., 2013) and establish sensitive biomarkers with parameters such as z-scores. However, challenges remain for accurate and reliable parcellation of specific group of human brain FNs especially neonate brain FNs with relatively low image quality (e.g. low signal-to-noise ratio (SNR)) and no functional atlas as spatial constraints.

Brain parcellation based on functional connectivity can be established by measuring similarity of time courses of rs-fMRI BOLD signals and aggregating brain voxels with similar BOLD signal time course pattern into clusters. A considerable number of algorithms have been developed for brain parcellation, including decomposition methods such as independent components analysis (ICA) (e.g. Calhoun et al., 2001; Beckmann et al., 2005; Damoiseaux et al., 2006) and non-negative matrix factorization (NNMF) (e.g. Anderson et al., 2014; Sotiras et al., 2015; Li et al., 2017), K-means (e.g. Kim et al., 2010; Kahnt et al., 2012; Chang et al., 2013), fuzzy k-means (e.g. Cauda et al., 2010, 2011), hierarchical clustering (Mumford et al., 2010; Bellect et al., 2006), spectral clustering (e.g. Van den Heuvel et al., 2008; Shen et al., 2010, 2013; Craddock et al., 2012; Cheng et al., 2014; Parisot et al., 2016; Wang et al., 2016; Shi et al., 2017), Gaussian Mixture Models (e.g. Yeo et al., 2011) and region growing (e.g. Heller et al., 2006). Other modality of neuroimaging data such as diffusion MRI has also been used to parcellate the brain (Glasser et al, 2016; Fan et al., 2016). The parcellations across various methods (e.g. Power et al., 2011; Yeo et al., 2011; Gordon et al., 2017) have been comparable because of unified underlying human brain organization. Many existing methods are sensitive to noise and may result in individual parcellated network lacking homogeneity and having spurious small patches. For example, the results of ICA-based methods (e.g. Calhoun et al., 2001; Beckmann et al., 2005) typically generate a large number of components in brain networks, the specific functions of which may be difficult to identify or classify. The K-means (e.g. Kim et al., 2010) and fuzzy-k-means (e.g. Cauda et al., 2010) methods can be fast and easily implemented, but are sensitive to noise or outliers of the fMRI BOLD signals. Recently developed parcellation methods based on normalized cut (Ncut) (Shi and Malik, 2000) has the advantages of being implemented relatively easily for brain parcellations (e.g. Van Den Heuvel et al., 2008; Shen et al., 2010; Craddock et al., 2012; Shi et al., 2017) and is robust to outliers. However, Ncut is sensitive to noise. It is difficult to remove the spurious patches using Ncut without imposing the constraints from the atlas (e.g. Shi et al., 2017) or spatial constraints (e.g. Craddock et al., 2012) that may disrupt the shapes of underlying networks. These Ncut-based parcellation methods are powerful tools to generate consistent regions of interests (ROIs) across individual subjects, but are not suitable for data-driven exploratory investigation such as studying early brain architecture with rs-fMRI data from neonate brains. The early brain development exhibits significantly changing structural (e.g. Huang and Vasung, 2014; Yu et al., 2016, Song et al., 2017; Ouyang et al., 2019) and functional (e.g. Doria et al., 2010; Cao et al., 2017a; Cao et al., 2017b) organizations that are different from adults. Lower intra-network connectivity of the higher-order FNs in the neonate brain presents challenges for the parcellation, as these higher-order FNs have not emerged (e.g. Doria et al., 2010; Cao et al., 2017a; Cao et al., 2017b). The lower signal-to-noise ratio (SNR) of neonate rs-fMRI data compared to that of adult brain data adds additional difficulty.

To meet these challenges, in this study we developed a novel data-driven Regularized-Ncut (RNcut) parcellation method that is robust to noise, generates smooth parcellated networks and significantly reduces the spurious small patches within each region. RNcut was tailored for noisy datasets (e.g. neonate brain datasets) and exploratory investigation with no functional atlas to be applied as spatial constraints. RNcut was established by adding two regularization terms to the conventional Ncut method. It was tested with simulated data with known ground truth and then applied to rs-fMRI of both adult and neonate brains. The RNcut software package is freely downloadable in the website www.nitrc.org/projects/rncut.

## Methods

Overview of the RNcut framework is illustrated in Fig 1. Time-series of rs-fMRI BOLD signal from each subject underwent preprocessing pipeline and were registered to the template space. Pearson’s correlation was then used to quantify similarity between all pairs of voxels (e.g. voxel *i* and voxel *j* in Fig 1) within the gray matter mask in each brain, resulting in the weight matrix for each subject. Mean weight matrix was obtained by averaging all individual weight matrices to represent the group pattern of functional connectivity. The RNcut method, highlighted by yellow, was applied to mean weight matrix to generate functional parcellation of the brain.

**Figure 1:**
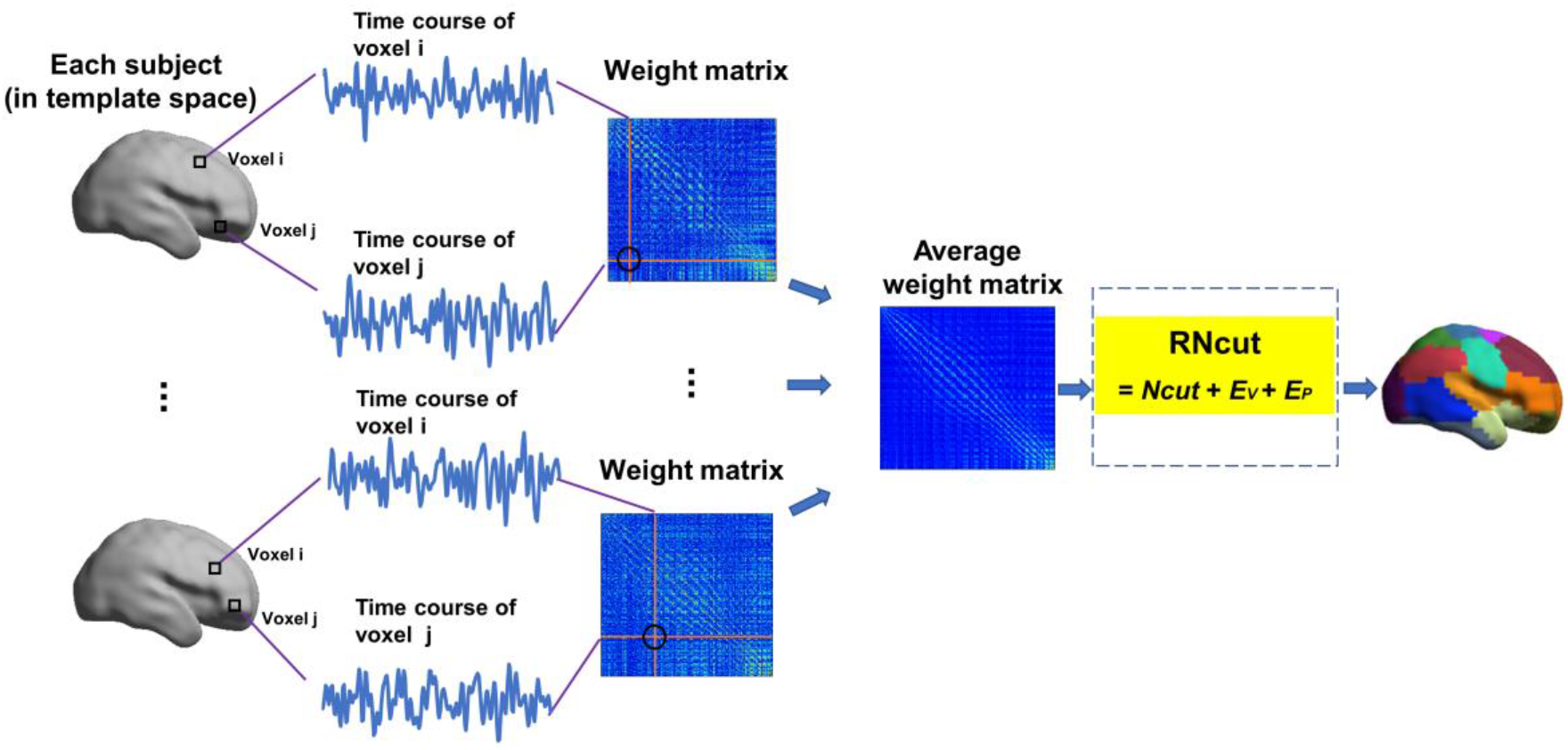
Flowchart of the Regularized-Ncut (RNcut) parcellation method which is composed of conventional Ncut and two regularization terms, 𝐸_𝑉_ (smoothing term) and 𝐸_𝑃_ (small-patch removal term).

### Brief description of normalized-cut (Ncut)

Given a rs-fMRI dataset, a graph *G=*{*V, E*} can be constructed, where *V* denotes all voxels in the cerebral cortex of the brain, and *E* represents connectivity strength quantified by weight *w*(*i,j*) usually calculated by Pearson’s correlation coefficient between the BOLD signals at voxel *v_i_* and voxel *v_j_.* Each brain voxel is a vertex of the graph *G*. The graph *G* can be cut into two disjoint sets *A*_1_ and *A*_2_. The cut cost was defined as the sum of the weights on edges connecting voxels in *A*_1_ to voxels in *A*_2_:

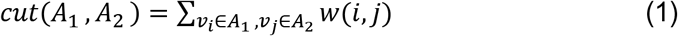

The optimal partition of a graph is the one that minimizes this cut cost. To avoid unnatural bias for partitioning out small sets of points, normalized-cut (Ncut) algorithm (Shi and Malik, 2000) was proposed. Ncut partitions the graph to minimize the disassociation measure between the groups, defined as:

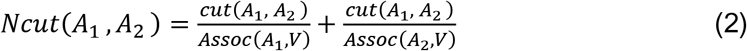

where *Assoc* 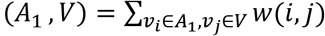 is the sum of weights on the edges between *A*_1_ to all voxels in the graph; similarly, *Assoc* 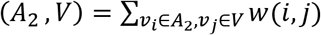 is the sum of weights on the edges between *A*_2_ to all voxels in the graph. The objective function of Ncut in Eq. 2 is updated in an iterative way until the difference between labels of vertices in next iteration and those in current one is less than a certain threshold (Shi and Malik, 2000).

For a given rs-fMRI dataset, time series of each voxel in the cerebral cortex was normalized to have zero mean and unit length. The weight matrix of individual subject *W_subject_* was calculated by Pearson correlation coefficient between each pair of voxel (*v_i_*, *v_j_*):

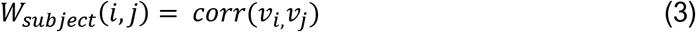

After constructing weight matrix for each subject, the group mean weight matrix *W* was obtained by averaging individual weight matrices. This group mean weight matrix was used for cut and Ncut in the equations (1) and (2), respectively.

### Regularized*-Ncut (RNcut)*

Since Ncut partitions the graph into different parts without considering underlying local structure of the graph, no spatial information about each voxel’s neighborhood structure was included as a regularization factor. To obtain spatially contiguous regions, it is common to explicitly introduce spatial neighborhood information for functional clustering of rs-fMRI data, typically by restricting the graph-edges to pairs of spatially neighboring voxels. For example, the previous work of whole brain parcellation using Ncut (Craddock et al., 2012) defined similarity between two voxels using a functional similarity measure with their spatial distance empirically specified. The whole brain was then parcellated into a large number of ball-shaped patches. Distance as a strict constraint may not be suitable for brain network parcellation since some networks (e.g., the default mode network) consist of several disjoint regions. Markov Random Field (MRF) defined over a graph (Tang et al., 2016; Kohli et al., 2009) and balanced by a tuning parameter *r* was used to cluster spatially contiguous voxels into the network while *r* was adjusted so that disjoint regions could be clustered into the same network. Weighted MRF was one regularization adopted in the RNcut. In addition, Ncut may result in small patches in certain sub-graphs due to noise. Spurious small patches usually generated due to noise need to be removed from the parcellated brain networks. A small-patch-removal term as another regularization was then adopted in the RNcut. The added two regularizations make RNcut method more robust to noise. Specifically, the presented RNcut method was established by adding the smoothing term (represented by MRF) and small-patch-removal term to Ncut, with its diagram shown in Fig. 2. RNcut was formulated in the equation below:

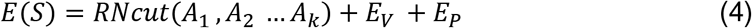

**Figure 2:**
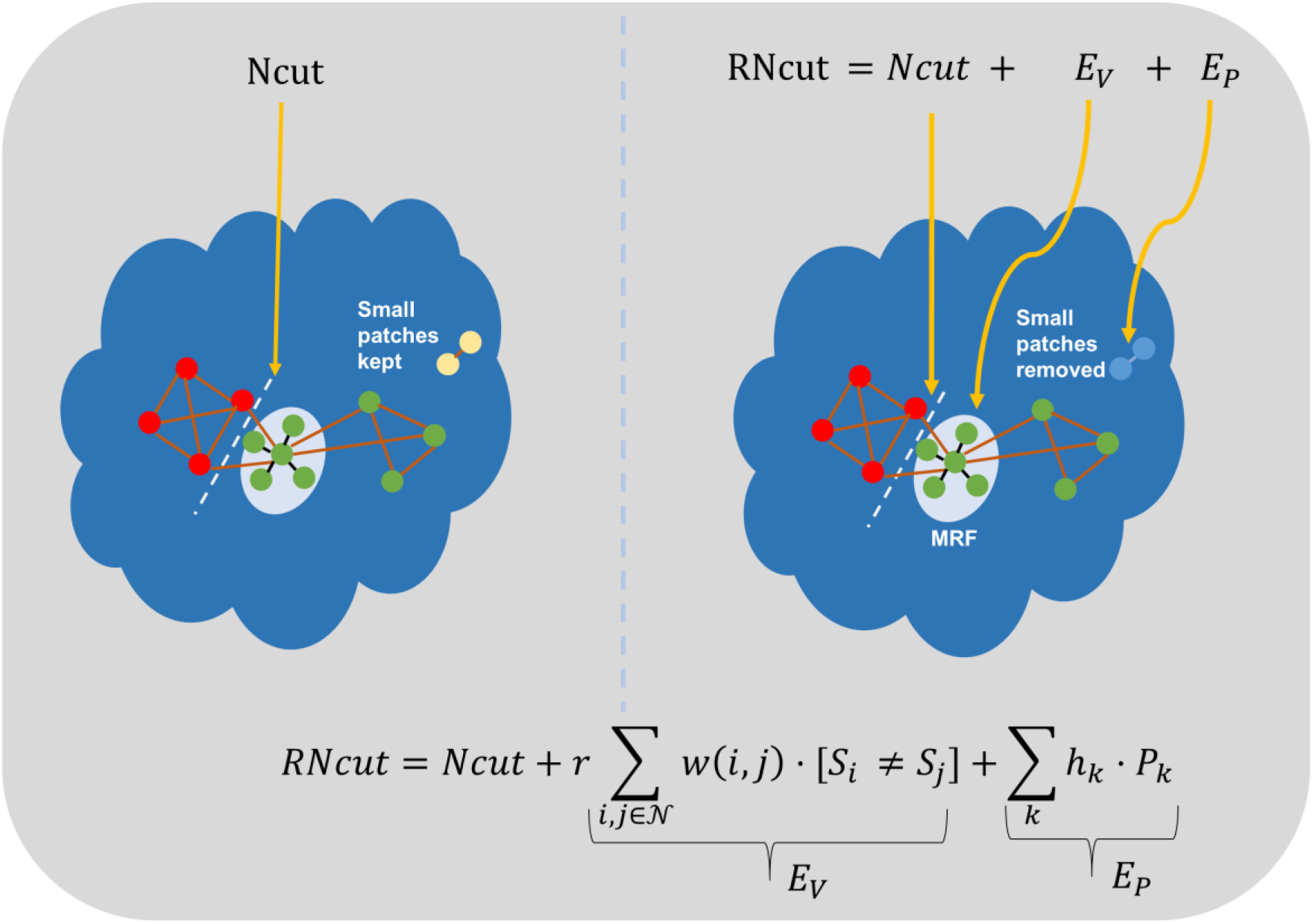
The diagram of the RNcut method. The left and right panel demonstrates how Ncut (see Eq. 2) and RNcut (see Eqs. 4-6) works, respectively.

where *A_k_* denotes the *k* different sub-graphs or networks; and *E_V_* and *E_P_* are the two regularization terms further formulated below. *E_V_* was defined by using MRF regularization (Tang et al., 2016):

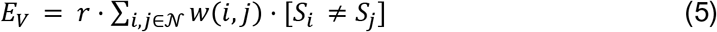

where 𝐸_𝑉_ is a smoothing term denoting the total penalty for discontinuity between voxel 𝑣_𝑖_ and voxel 𝑣_𝑗_, which are neighborhood voxels; *S_i_* and *S_j_* are labels for voxel *i* and voxel *j*; and *r* is a tuning parameter. A small-patch-removal term *E_P_* was formulated as:

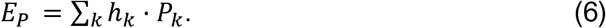

where *h_k_* = log(*P_k_*) is the penalty cost for each brain network, and *P_k_* is the number of sub-networks (or patches) in the *k*th network. Each brain network consists of limited number of patches. By adding the penalty cost to large number of patches in each network, it will suppress spurious small patches and obtain the brain networks robust to noise.

As the objective Eq. 4 can be minimized by the alpha-expansion method (Boykov et al., 2001), an iterative strategy (Wang et al, 2015) was utilized to perform the group parcellation with the algorithm described below.

#### Algorithm 1 (RNcut parcellation)

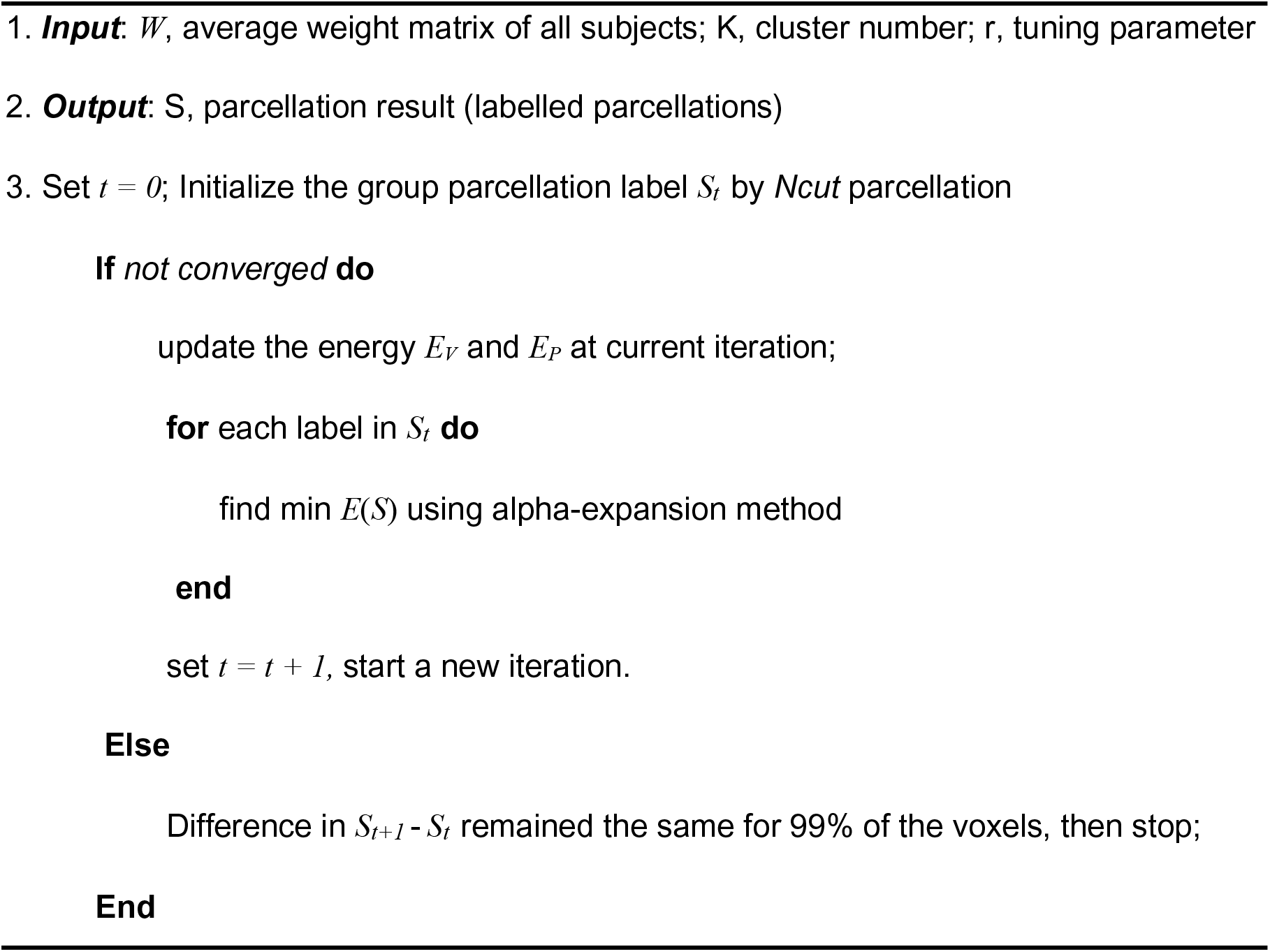

### Rs-fMRI datasets and preprocessing

#### Simulated datasets

The simulated fMRI dataset was generated using the SimTB toolbox (Erhardt et al. 2012) which allows flexible generation of time-series of fMRI signals (http://mialab.mrn.org/software/simtb/). The dataset consisted of simulated data of 20 subjects. Simulated dataset of each subject included 150 2D images with dimension 100×100 and 15 distinctive FNs. Rician noise was added to the simulated images and contrast-to-noise ratios (CNR) of the noisy image ranged from 0.1 to 0.3.

#### Adult datasets and preprocessing

Two groups of adult brain rs-fMRI datasets with different dynamics, one with dynamics of 225 and the other with dynamics of 128, were used to evaluate how the proposed RNcut performs with long and short time series of rs-fMRI datasets. Both groups of adult datasets (Beijing Zang and ICBM) were downloaded from a public website www.nitrc.org/projects/fcon_1000 (Biswal et al. 2010). The adult brain group #1 datasets (Beijing Zang) included those from 26 subjects. A total of 225 whole brain echo-planar imaging (EPI) volumes were acquired with a T2-weighted gradient-echo sequence using the following parameters: TR = 2s, in-plane imaging resolution = 3.125 × 3.125 mm^2^, in-plane field of view (FOV) = 200 × 200 mm^2^, slice thickness = 3.6 mm with no gap; slice number = 33. Before preprocessing, the first 10 volumes were removed for rs-fMRI signal to reach a steady state so that 215 functional volumes was kept for each subject. These rs-fMRI datasets were corrected for acquisition time delay between slices and head motion between volumes. They were then registered to the template space. The normalized image volumes were further filtered by a Gaussian kernel with full width at half-maximum of 6 mm, and underwent linear trend removal and temporal band-pass filtering (0.01-0.10Hz). Finally, several nuisance variables, including head motion parameters and averaged signal from white matter and cerebrospinal fluid tissue, were regressed out to reduce the effects of non-neuronal signals. The above-mentioned procedures (Calhoun et al., 2017) were conducted by using DPABI (http://rfmri.org/dpabi, Yan et al., 2016). The same procedures were also conducted on the adult brain group #2 datasets (ICBM) from 31 subjects with rs-fMRI dynamics of 128.

#### Neonate dataset and preprocessing

A series of MR sequences including rs-fMRI were conducted on neonates with a Philips 3T Achieva MR scanner (Yu et al., 2016; Cao et al., 2017a; Ouyang et al., 2017; Song et al., 2017). Rs-fMRI datasets of 19 term-born neonates (37-41.7 postmenstrual weeks) with head motions below the threshold (see Cao et al., 2017a for details of head motion analysis) were used in this study. All neonate rs-fMRI datasets can be freely downloaded from a website maintained by us (www.brainmrimap.org). All MR scans were performed during natural sleep of neonates without sedation. A written consent approved by Institutional Review Board (IRB) was obtained from the parent for every participating neonate. A T2-weighted gradient-echo EPI sequence was used to acquire the rs-fMRI. A total of 210 whole brain EPI volumes were acquired using the following parameters: TR = 1500 ms, TE = 27 ms, flip angle = 80°, in-plane imaging resolution = 2.4 × 2.4 mm^2^, in-plane FOV = 168 × 168 mm^2^, slice thickness = 3 mm with no gap, slice number = 30. The rs-fMRI scan time was 5.4 minutes. A co-registered T2-weighted image was acquired with turbo spin echo (TSE) sequence as the structural MRI with the following parameters: TR = 3000 ms, effective TE = 80 ms, in-plane imaging resolution = 1.5 × 1.5 mm^2^, in-plane FOV = 168 × 168 mm^2^, slice thickness = 1.6 mm with no gap, slice number = 65. Before preprocessing, the first 15 volumes of rs-fMRI dataset were removed for the signal to reach a steady state and remaining 195 functional volumes were retained for each subject. After correction for acquisition time delay between slices and head motion between volumes, the rs-fMRI image volumes were then registered to the custom-made template (Cao et al., 2017a). The same filtering and regression preprocessing procedures described in details in our previous publication (Cao et al., 2017a) were applied.

### Quantitative evaluation and comparison of different parcellation methods

Competing methods including ICA, NNMF, Ncut and spatially constrained spectral clustering (SCSC) (Craddock et al., 2012) were used as the references for evaluating the performance of RNcut. Group ICA of fMRI Toolbox (GIFT) (Calhoun et al., 2001).was adopted for decomposing the fMRI data from a group of subjects for ICA method. The mean weight matrix described was used for parcellating the simulated data for three other methods, NNMF (Anderson et al. 2014), Ncut (Shi et al., 2000) and the RNcut. Different parcellation methods were quantitatively evaluated and compared with both simulated and human subject datasets. The following evaluation measures were used: (1) Dice coefficient (Dice, 1945) and L1 error (Ratnanather et al., 2004), (2) Silhouette Index (SI) (Rousseeuw, 1987) for measuring parcellation homogeneity, and (3) number of patches in each network.

#### Dice coefficient

Dice coefficient quantifying overlap of two labelled regions (region A and B) was obtained with the equation:

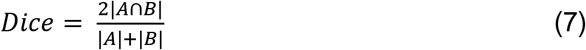

where | · | indicates the number of vertices in a region.

#### L1 error

The L1_error was defined as:

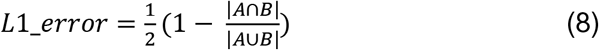

Larger value of the Dice coefficient and smaller value of the L1_error indicate higher parcellation accuracy.

Homogeneity measurement: Silhouette index (SI) was used for quantifying the intra-network (or intra-class) homogeneity. SI normalizes *a_k_* by the maximum similarity between every cluster voxel and every out-of-cluster voxel (Craddock et al., 2012). Similar to the homogeneity measures used in the literature. SI was calculated with the following equations

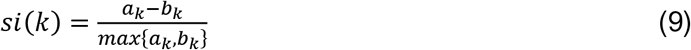

where

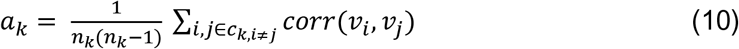

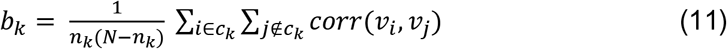

In Eqs. 10 and 11, *ck* denotes *k*th network and *nk* the number of voxels in this network; 𝑁 is the total number of voxels and SI is the averaged *si(k)* of all networks. The SI value of 1 indicates perfect homogeneity.

#### Number of patches in each network

It is a measure to quantify the fragmentation of each network. The larger value indicates that the network has more isolated patches, many of which are usually spurious small patches caused by noise.

The pair-wise quantitative comparisons of these evaluation measures between different parcellation methods were conducted with t tests. The quantitative comparisons of intra-network connectivity of parcellated network between neonates and adults and quantitative comparisons between intra- and inter-network connectivity of neonates were also conducted with t tests.

### Reproducibility tests of RNcut

Reproducibility of the proposed RNcut was tested using the adult and neonate datasets described above. Specifically, for adult brains, RNcut was applied to each of the two non-overlapping subgroups obtained by evenly dividing the total 26 subjects from the datasets of adult brain group #1. For neonate brains, the group of total 19 neonate subjects were divided into two non-overlapping subgroups with 10 and 9 neonate subjects in each subgroup, respectively. RNcut was applied to each of the two subgroups. Dice coefficients were calculated to quantify the reproducibility.

## Results

### Parcellation results of simulated datasets using RNcut compared to those using ICA, NNMF, Ncut and SCSC

The FN number 15 was set for all methods for comparison. Fig. 3 shows the parcellation results using ICA, NNMF, Ncut, SCSC and RNcut methods on the simulated data added with noise to test robustness of each method against noise. The CNR of the tested noisy simulated data is 0.1-0.3. The FNs from different methods were evaluated using the Dice coefficient and L1_error. Fig 3 demonstrates that overall RNcut outperformed ICA, NNMF, Ncut and SCSC. Specifically, Fig 3a shows that RNcut is more robust to noise with almost no spurious voxels while parcellation results from ICA, NNMF and Ncut were significantly affected by noise, indicated by widespread spurious voxels. Quantified measurements, including Dice coefficient in Fig. 3b and L1 error in Fig. 3c, further confirmed that RNcut is more robust against noise compared to the ICA, NNMF, SCSC and Ncut. The measured numbers of spurious pixels in all clusters in Fig. 3d indicate that RNcut and SCSC yield much smaller number of spurious pixels than ICA, NNMF and Ncut. Despite yielding fewer spurious pixels, SCSC disrupts the shapes of the parcellated regions (Fig. 3a) due to the constraints of equal sizes of the parcellated regions. For example, the three regions with different colors on the bottom of ground-truth image (middle left panel of Fig. 3a) were mistakenly integrated into one in parcellations using SCSC (middle right panel of Fig. 3a), pointed by the yellow arrows.

**Figure 3:**
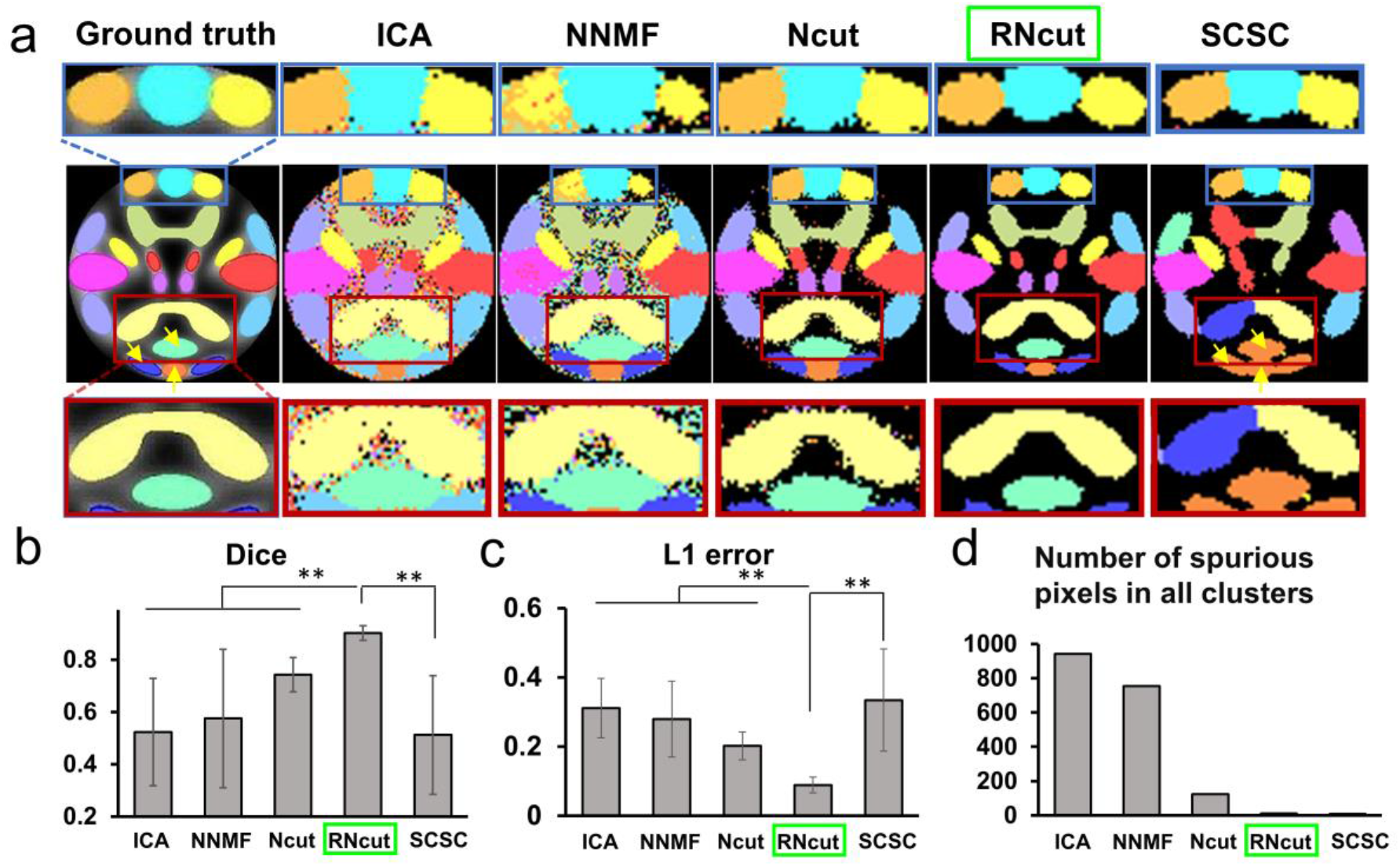
Parcellation of 15 networks with simulated data using ICA, NNMF, Ncut, RNcut and SCSC as well as quantitative comparisons of the parcellation performance of these methods. In panel a, noise was added to simulated dataset, resulting in CNR ranged in 0.1-0.3. The regions marked by red rectangle showed the proposed RNcut could generate clean and smooth clusters. The regions marked by blue showed the proposed RNcut could generate better small clusters closer to ground truth and with much smaller number of spurious pixels. The measurements of Dice ratio (b) and L1 error (c) as well as the images in panel a demonstrate that RNcut is more robust to noise than all other methods. The measurements of number of spurious pixels in all clusters (d) show that the RNcut outperformed the ICA, NNMF and Ncut while yielding similarly small number of spurious pixels to what SCSC yields. ** indicates p<0.01.

Measurements of the Dice coefficient and L1 error were used to optimize the tuning parameter *r* for the MRF regularization term *E_V_* in the Eq. 5. As shown in Fig 4, by changing different tuning parameter r in the Eq. 5, the highest Dice coefficient and lowest L1 error for simulated datasets were achieved with r =5. For adult and neonate brain datasets tested below, the value of r was tested in the range of 2 to 8. As shown in the Supplementary Fig 1, the SI is highest with r value set to 3 and 5 for the adult and neonate datasets, respectively. Thus r value of 3 and 5 was used in RNcut for brain parcellations with adult and neonate rs-fMRI below, respectively.

**Figure 4:**
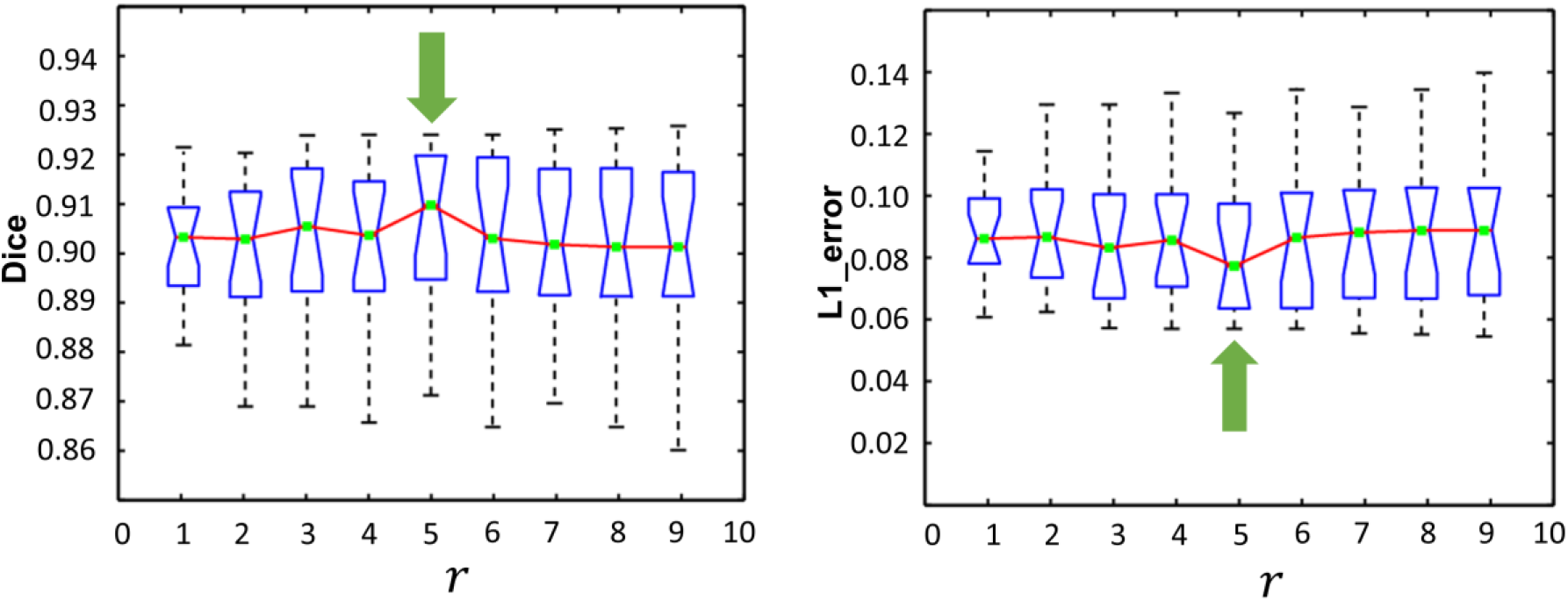
Selection of optimized tuning parameter r for MRF regularization. Dice ratio and L1 error vary with different r with the green arrow indicating optimized r associated with highest Dice ratio and lowest L1 error.

### Parcellation results of the adult and neonate brains using RNcut compared to those using ICA, NNMF, Ncut and SCSC

The cortical parcellation was evaluated on the human subject datasets including two groups of adult datasets and one group of neonate datasets. 17 FNs have been consistently identified in the human brain (Yeo et al. 2011; Gordon et al., 2014). Therefore, the network number 17 was set for functional parcellations of the adult and neonate brains. Fig. 5a shows the FNs after ICA, NNMF, Ncut, SCSC and RNcut were applied to the first group of adult brain datasets including rs-fMRI datasets of 26 adult subjects with dynamics of 225. It can be appreciated from Fig. 5a that much fewer spurious patches in parcellated FNs, indicated by red arrows, were created by the RNcut method. Fig. 5b shows that statistically significantly higher SI value was obtained with the parcellations from RNcut, compared to parcellations from ICA, NNMF, SCSC and Ncut. Fig. 5c demonstrates that smaller number of isolated patches in the parcellated networks were generated by the RNcut compared to ICA, NNMF and Ncut. SCSC yielded even smaller number of isolated patches (Fig. 5c), but disrupted the shapes of underlying adult brain functional networks (Fig. 5a). Higher homogeneity is associated with RNcut demonstrated by Fig 5b. ICA, NNMF, Ncut, SCSC and RNcut were also applied to the second group of adult brain datasets including rs-fMRI datasets of 31 adult subjects with shorter dynamics of 128. Significantly better performance of the parcellations was obtained with RNcut too, compared to other competing methods, as shown in the Supplementary Fig 2.

**Figure 5:**
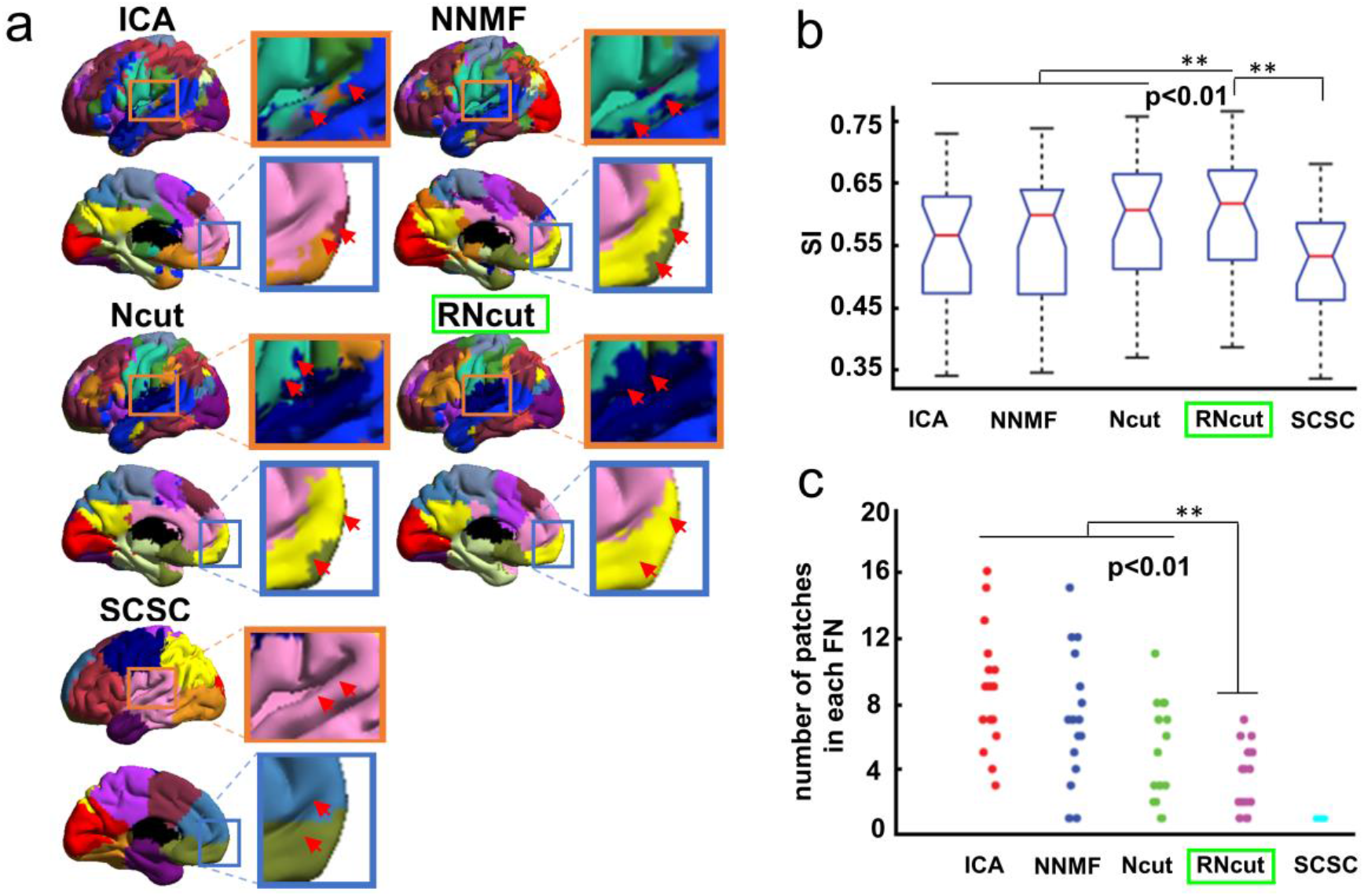
Parcellation of the functional networks of adult brains using different methods. In panel a, the parcellation results of ICA, NNMF, Ncut, SCSC and RNcut demonstrate that fewer isolated spurious patches were obtained with RNcut, indicated by red arrows and highlighted with the enlarged boxes. Panel b shows significant higher Silhouette Index (SI) value of RNcut method than all other competing methods. Panel c shows significantly smaller number of spurious patches in each functional network generated by RNcut method, compared to ICA, NNMF and Ncut. ** indicates p<0.01.

Besides adult brain datasets, ICA, NNMF, Ncut, SCSC and RNcut were applied to the neonate dataset. From Fig 6a, isolated spurious patches, indicated by red arrows, are more prominent in the parcellated networks generated with ICA, NNMF and Ncut. In contrast, the FNs generated with RNcut include much fewer spurious patches. Fig. 6b shows that statistically significantly higher SI value was obtained with the parcellations from RNcut, compared to parcellations from ICA, NNMF, SCSC and Ncut. Fig. 6c demonstrates that RNcut generated smallest number of isolated patches in the parcellated neonate brain networks, compared to ICA, NNMF and Ncut. Similar to Fig. 4 and Fig. 5, SCSC was able to yield even smaller number of isolated patches (Fig. 6c), but disrupted the shapes of underlying neonate brain functional networks (Fig. 6a).

**Figure 6:**
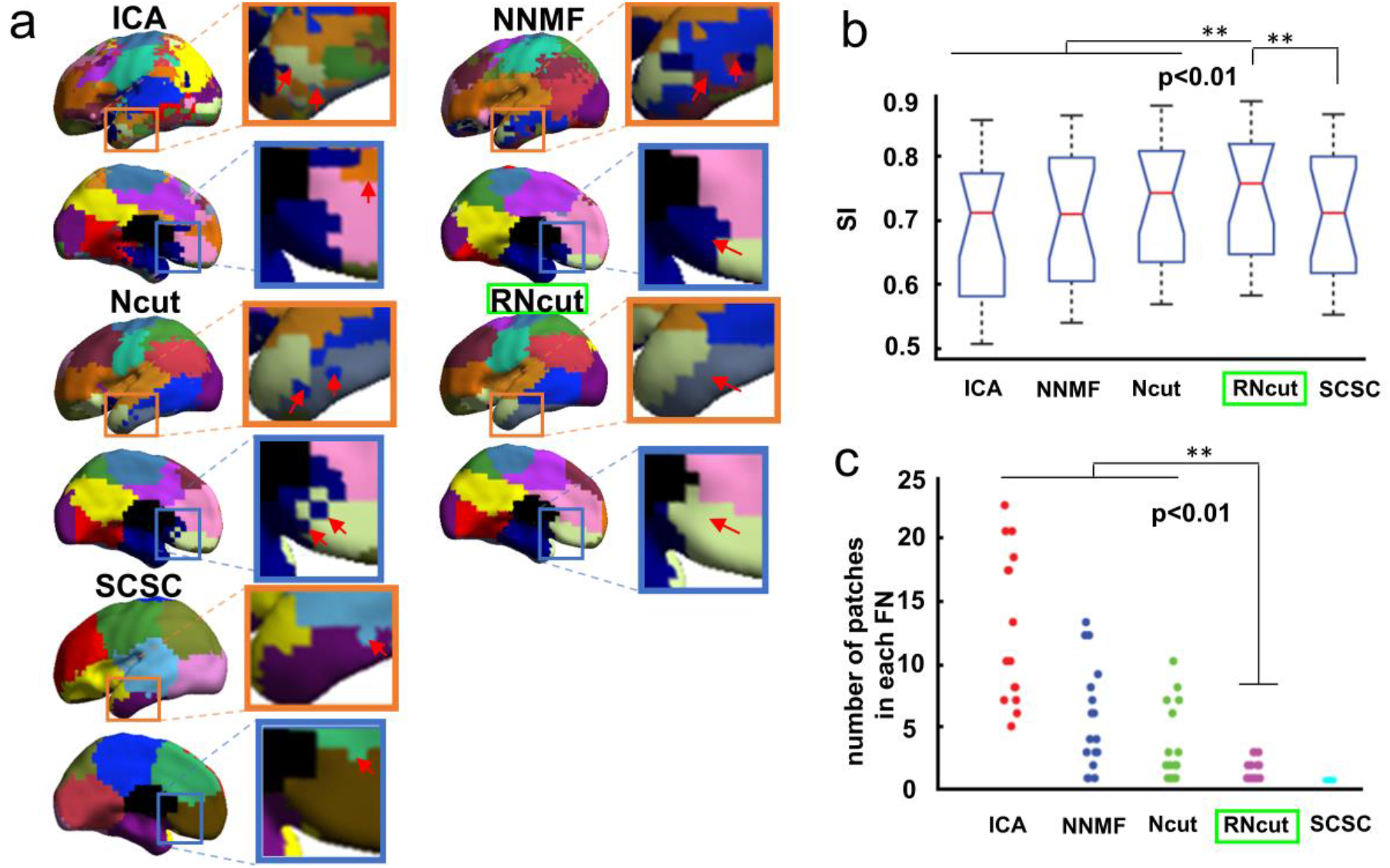
Parcellation of the functional networks of neonate brains using different methods. In panel a, the parcellation results of ICA, NNMF, Ncut, SCSC and RNcut demonstrate that fewer isolated spurious patches were obtained with RNcut, indicated by red arrows and highlighted with the enlarged boxes. Panel b shows significant higher Silhouette Index (SI) value of RNcut method than all other competing methods. Panel c shows significantly smaller number of spurious patches in each functional network generated by RNcut method, compared to ICA, NNMF and Ncut. ** indicates p<0.01.

### Reproducibility of parcellation using RNcut

The parcellation results of 2 adult brain subgroups (Fig. 7a) and 2 neonate brain subgroups (Fig. 7b) demonstrate that the RNcut parcellations are highly reproducible for both adult and neonate brains. The high reproducibility is also indicated by the high Dice ratio shown in Fig. 7c. Among adult brain parcellated networks, reproducibility of parcellations of auditory, visual and default-mode networks, indicated by red arrows in Fig 7a and reflected by higher Dice ratio, is higher than that of other networks. Reproducibility of inferior parietal and inferior temporal networks in the adult brains, indicated by black arrows in Fig. 7a and reflected by lower Dice ratio, is lower than that of other networks. Among neonate brain parcellated networks, reproducibility of parcellations of visual and lateral parietal networks, indicated by red arrows in Fig 7b and reflected by higher Dice ratio, is higher than that of other networks. Reproducibility of ventral attention network and fronto-parietal networks in the neonate brains, indicated by black arrows in Fig. 7b and reflected by lower Dice ratio, is lower than that of other networks.

**Figure 7:**
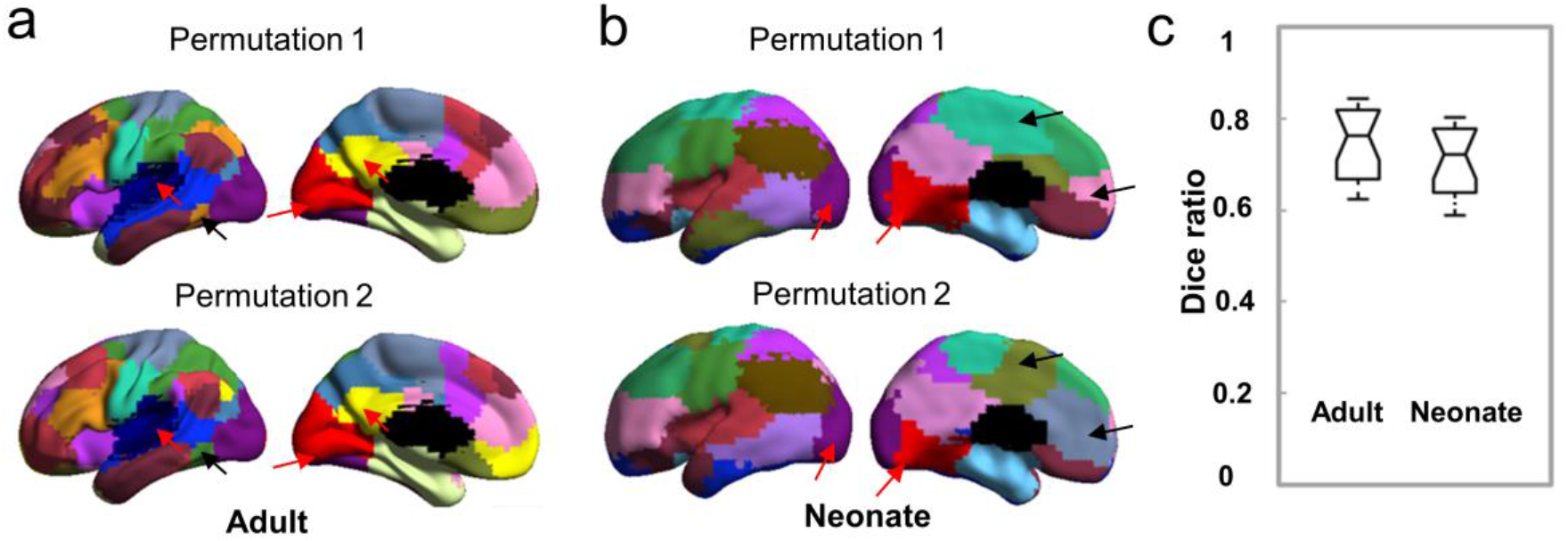
Reproducible parcellation by RNcut for adult and neonate brains. Panel a shows the parcellation results of two independent subgroup datasets from the adult group #1. Panel b shows the parcellation results of two independent subgroup datasets from the neonate group. Panel c shows the Dice ratio of the two parcellation results within adult or neonate group.

### Intra-network connectivity measured based on RNcut parcellation for both neonate and adult brains

Based on FNs parcellated by RNcut, the calculated intra-network functional connectivity indicates distinctive pattern in neonate brain compared to adult brain. The intra-network connectivity of each voxel was calculated as averaged Pearson correlation coefficient between this voxel and all other voxels in the same network. Fig. 8a and Fig. 8b show the intra-network connectivity map for neonate and adult brain based on FNs parcellated by RNcut. In the neonate brain (Fig. 8a), relatively higher intra-network connectivity was only found in the dorsal and ventral sensorimotor (DSM and VSM) network. On the contrary, relatively higher intra-network connectivity was much more widespread in the adult brain (Fig. 8b) covering almost all major networks as these networks have become fully mature in adults. With Fig. 8c demonstrating quantitative comparison of intra-network connectivity between neonates and adults in different FNs, it is intriguing to observe that the only network with no significant difference of intra-network connectivity strength between the neonate and adult brain is the primary sensorimotor network (DSM and VSM). Intra-network connectivity in all other networks of neonate brain is significantly lower than that of the adult brain. As shown in Fig. 8d, exclusively higher intra-than inter-network connectivity in all networks was found for neonate brain, possibly due to dominant short-range connections in the neonate brain. Compared to those generated with FNs parcellated by ICA, NNMF or Ncut, intra-network connectivity maps generated with RNcut displayed a clear and unique pattern in neonate brain, as shown in Supplementary Fig. 3. Specifically, the high contrast aligned with the boundary of primary sensorimotor area (DSM and VSM) was shown in the intra-network connectivity maps generated with RNcut.

**Figure 8:**
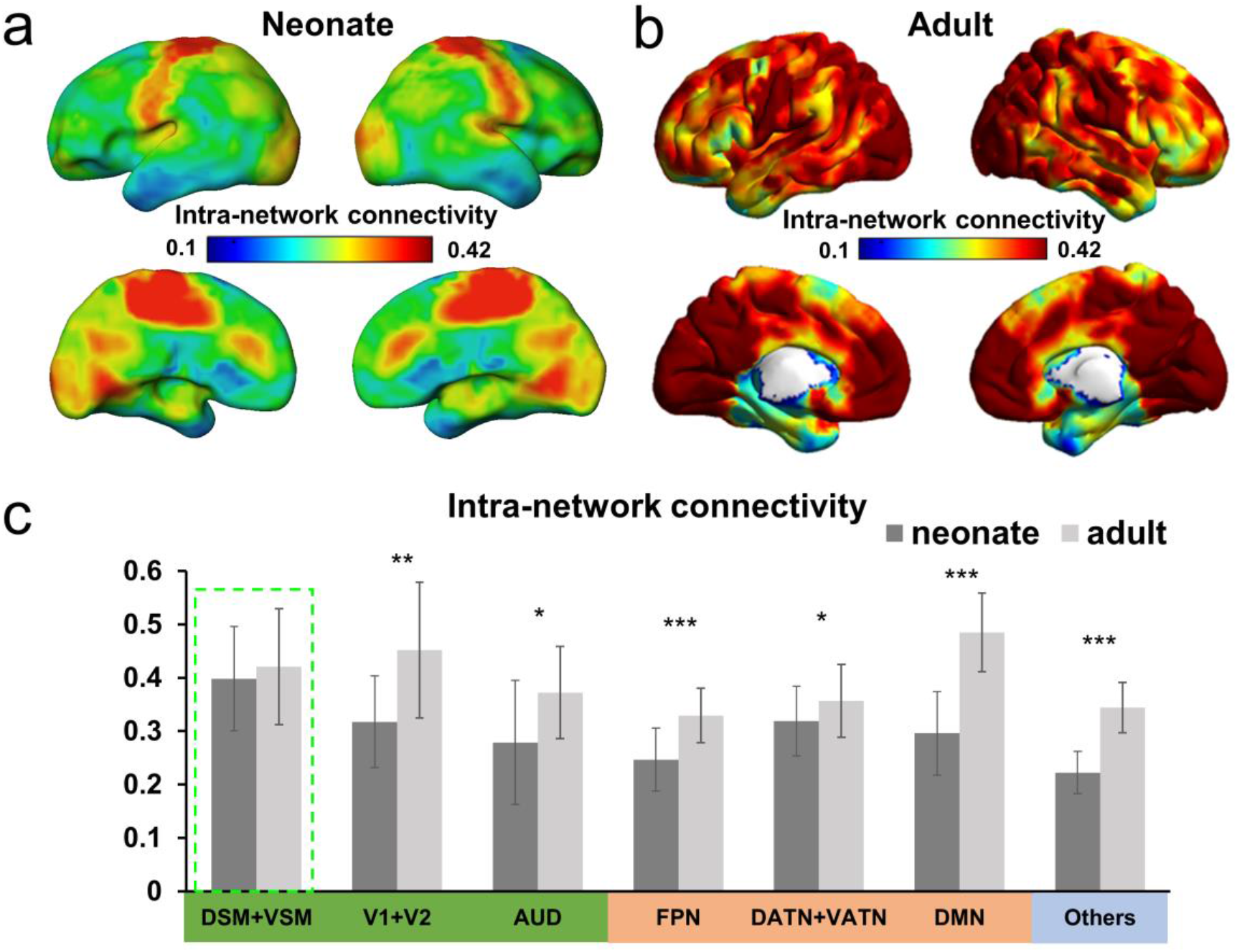
Intra-network connectivity for neonate and adult brain based on parcellated networks with RNcut. Panel a and b show the intra-network connectivity maps of neonate and adult brain, respectively; Panel c shows that except dorsal and ventral sensorimotor networks (DSM and VSM in green row), intra-network connectivity in all other networks (pink and blue row) of neonate brain is significantly lower than that of adult brain,. Panel d shows that the intra-network connectivity is exclusively higher than inter-network connectivity in all networks of the neonate brain. Abbreviations: V1+V2: Primary and secondary visual; AUD: Auditory; FPN: Fronto-parietal network; DATN+VATN: Dorsal and ventral attention network; DMN: Default mode network. * indicates p<0.05; ** indicates p<0.01; *** indicates p<0.001.

## Discussion

In this study, we presented a novel data-driven and connectivity-based algorithm called RNcut. RNcut was established by integrating conventional Ncut with two regularization terms to generate robust and homogeneous functional parcellations of both adult and neonate human brains. The RNcut has several significant improvements compared to conventional Ncut in that (1) RNcut is robust to noise and capable of parcellating functional regions reproducibly and more accurately, especially those regions with relatively weak intra-network functional connectivity; (2) Most of small patches representing spurious FNs can be identified and eliminated by RNcut. Consequently, functional regions with higher spatial contiguity are generated by RNcut parcellation; (3) High functional homogeneity within each parcellated FN can be achieved with RNcut; and (4) RNcut can be applied to not only adult brains but also neonate brains characterized with unique rs-fMRI signal patterns. RNcut is especially suitable for exploratory investigation with noisy datasets (e.g. neonate data) to facilitate addressing a suite of clinical and neuroscientific questions related to maturational patterns of the FNs.

The improvements of RNcut were supported by the results obtained from simulated, adult and neonate brain rs-fMRI datasets. Test results from the simulated datasets demonstrated that RNcut parcellation was robust to noise (Fig. 3). With optimized tuning parameter *r* for the RNcut algorithm (Fig. 4), homogeneous parcellations of adult (Fig. 5) and neonate (Fig. 6) brains were observed. RNcut outperformed competing methods including ICA, NNMF, SCSC and Ncut (Fig. 3, 5 and 6) and showed high reproducibility in both adult and neonate brains (Fig. 7). Intra-network connectivity was quantified in neonate and human brains with the parcellated networks by RNcut, revealing distinctive intra-network connectivity patterns in the neonate brains compared to those in the adult brains (Fig. 8). These improvements included good spatial contiguity with relatively fewer spurious patches, higher functional homogeneity in each parcellated network and reproducible parcellations of noisy datasets. The improvements were achieved by two added regularization terms. The first regularization term leverages the pattern of neighboring nodes that have high possibility of sharing same label in a graph. This regularization was realized by using the Markov Random Field (MRF) (Tang et al., 2016; Kohli et al., 2009). The combination of Ncut and MRF so far has been only used to process static images (Tang et al., 2016), not to parcellate the brain through processing BOLD signal time courses in rs-fMRI data. By considering neighborhood information, MRF can smooth the local boundaries. The second regularization term was originally brought in this study and imposed a penalty on small spurious patches in each network. To our knowledge, RNcut algorithm is the first incorporating these two regularization terms for functional brain parcellation. Other Ncut-based methods focused on consistency of the ROIs across the individual subjects at the cost of disrupting the underlying FN shapes and decreasing the homogeneity of each FN. For example, the structural constraints in the previous work (Carddock et al. 2012) tend to generate sphere-like parcellated regions with similar sizes and large parcellation numbers. Compared to these methods, RNcut has the advantage of achieving spatial contiguity and eliminating the spurious patches with dataset of lower SNR without imposing strong spatial constraints and without significantly affecting the underlying shapes of functional RNs. It thus can be used for exploratory data-driven investigations of the brain networks.

RNcut generates FNs with higher functional homogeneity compared to competing methods. Based on Hebb’s principle that “neurons that fire together wire together”, the rs-fMRI time courses of the voxels in the same intrinsic FN are likely to have higher correlations. Homogeneity within individual parcellated network, quantified by SI values, was used for evaluating the quality and accuracy of parcellations. Higher homogeneity inside each of the networks parcellated by RNcut resulted in higher SI values compared to those of the networks parcellated by ICA, NNMF, SCSC or Ncut. RNcut is able to generate robust group parcellation in not only adult, but also neonate brains. Most of existing methods (e.g. Calhoun et al., 2001; Beckmann et al., 2005; Van den Heuvel et al., 2008; Kim et al., 2010; Kahnt et al., 2012; Yeo et al., 2011; Chang et al., 2013; Anderson et al., 2014; Sotiras et al., 2015) demonstrated the parcellations of only adult brains. A recent pioneering study (Shi et al., 2017) was among the first to parcellate the FNs of infant brains. The parcellation method in this study was also based on Ncut and required spatial restriction from the automated anatomical labeling (AAL) atlas, resulting in relatively large number (around 90) of parcellated regions with relatively small sizes. Application of RNcut does not require information *a priori* and is able to generate networks with known functional meanings and larger sizes. The RNcut can then be potentially used to study early brain maturational patterns given heterogeneous and highly dynamic development of brain systems during this stage. The data-driven parcellation results from RNcut can potentially be used to establish the neonate- or infant-specific functional atlases. Distinctive intra-network connectivity pattern in neonate brain (Fig. 8) was revealed with RNcut parcellation and this pattern underlies heterogeneous emergence of brain functional systems during early development. Specifically, the intra-network connectivity of neonate brain (Fig. 8a) based on RNcut parcellation suggested that primary sensorimotor networks emerge earlier than other higher-order cognitive networks during early brain maturation, consistent with previous findings (e.g. Gao et al., 2009; Doria et al., 2010; Smyser et al., 2010; Cao et al, 2017a). Exclusively higher intra-than inter-network connectivity in all networks (Fig. 8d) obtained with RNcut parcellation may be due to dominant short-range connections in the neonate brain (Cao et al., 2017a). RNcut can also be readily applied to pediatric populations of other ages from birth to adulthood.

A few other technical considerations and limitations are elaborated below. First, since the presented RNcut method is based on Ncut, it is not readily applied to individual parcellations. One possible solution is to parcellate the individual brain with the guidance of the group parcellation and pre-estimated inter-subject variability map (e.g. Wang et al., 2015). Another possible solution is to add one more term representing variability information of individual subject to current RNcut algorithm (Eq. 4). Second, it is difficult to determine the tuning parameter in the proposed method. The optimal tuning parameter *r*=5 was obtained by quantifying the parcellation performance using the simulated datasets. The tuning parameter r should be adjusted based on the SNR of the rs-fMRI dataset to be processed. If the SNR of the rs-fMRI is low, it is recommended that a larger value of r is adopted to make the effects of neighboring voxels stronger (i.e. stronger MRF regularization) when determining the parcellation of the target voxel. The tuning parameter *r* with testing range value from 2 to 8 can be optimized with SI values obtained from parcellations of real human rs-fMRI data, as demonstrated in Supp Fig 1. Thirdly, selection of optimal number of brain parcellations remains to be an open problem for developing parcellation algorithms (e.g. Power et al., 2011; Yeo et al., 2011; Craddock et al., 2012; Thirion et al., 2014; Wang and Wang, 2016). Establishing an appropriate cost function integrating different measurements such as SI and Dunn index (Dunn, 1974) may shed light on determining optimal parcellation number. Unlike adult brain, little is known for neonate brain parcellations so far with many regions not well differentiated and characterized by clear functional signature (e.g. Doria et al, 2010; Cao et al., 2017a; Cao et al., 2017b; Ouyang et al, 2019). Therefore, only small number of parcellated networks was tested, resulting in relatively large area in each parcellated patch. The small parcellated patches were then considered as “spurious” ones with the assumption that it was not likely that a big patch and a small isolated one belong to the same functional area when the brain is highly immature. Identification of “spurious” patches is only possible after a better understanding of the “true” small patches. In the future, better understanding of the small patches could be obtained with lengthy and high-SNR rs-fMRI scans of many dynamics, a higher number of parcellated patches and reproducibility tests of the parcellated small patches across individual neonate subjects. The RNcut software package has been made publicly available in the website www.nitrc.org/projects/rncut for free use and further improvement.

## Conclusion

We proposed a novel RNcut algorithm for a robust parcellation of human brain cerebral cortex with rs-fMRI datasets. RNcut adopted two regularization terms to generate robust and homogeneous functional parcellation of not only adult but also neonate brain networks. Based on quantified evaluation results from both simulated and human subject datasets, RNcut outperformed widely used ICA, NNMF, SCSC and Ncut methods. RNcut is in general more robust to noise and generates parcellated networks with higher functional homogeneity, more smooth boundaries and less spurious small patches in each network. Intra-network connectivity quantified with RNcut parcellations revealed distinctive intra-network connectivity patterns in the neonate brains compared to those in the adult brains .RNcut could delineate the FNs across developmental ages and potentially facilitate the research on human brain functional development patterns.

## Acknowledgment

This work was supported by the National Institutes of Health (NIH) MH092535, MH092535-S1, HD086984 and EB022573.

## Supplementary material

Supplementary material associated with this article is provided.

