## Supplementary Information for "Regularized-Ncut: Robust and homogeneous functional parcellation of neonate and adult brain networks"

[1]Department of Radiology, Children's Hospital of Philadelphia, Philadelphia, PA, USA, [2]Department of Radiology, Perelman School of Medicine, University of Pennsylvania, Philadelphia, PA, USA, [3]Department of Bioengineering, School of Engineering and Applied Science, University of Pennsylvania, Philadelphia, PA, USA, [4]Department of Biomedical Engineering, School of Medicine, Tsinghua University, Beijing, China.

### *Supplementary Information*

Selection of optimal tuning parameter *r* for application of regularized normalized cut (RNcut) to adult and neonate brain datasets

Supp. Fig 1 shows the measurements of Silhouette index (SI) of the adult group #1 and neonate brain resting-state fMRI (rs-fMRI) datasets with varying tuning parameter r values from 2 to 8. Tuning parameter r represents the weight of regularization of Markov Random Field (MRF) in Eq. 5. The highest SI was associated with r=3 and r=5 for the adult and neonate brain datasets, respectively.

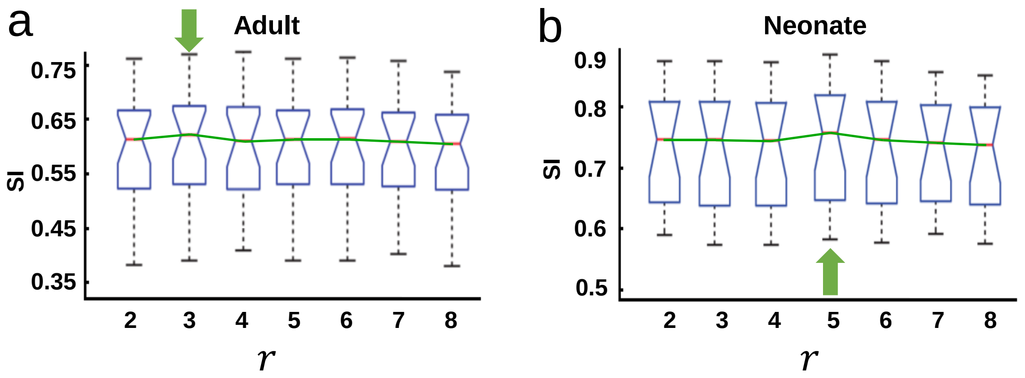

**Supplementary Figure 1: Panel a and b show the changes of SI measurements with varying tuning parameter r values from 2 to 8 for adult (a) and neonate (b) brain rs-fMRI datasets, respectively. Optimized r value 3 and 5, pointed by the green arrows, is associated with the highest SI measurement for adult (a) and neonate (b) dataset, respectively.**

Functional parcellation on second group of adult brain datasets

*Second group of adult brain datasets:* The adult group #2 datasets from 31 adult subjects were used to evaluate independent component analysis (ICA), non-negative matrix factorization (NNMF), normalized cut (Ncut), spatially constrained spectral clustering (SCSC) and regularized Ncut (RNcut). A T2-weighted gradient-echo echo-planar-imaging (EPI) sequence was used for the rs-fMRI scan. A shorter dynamics of 128 whole brain EPI volumes were acquired using the following parameters: TR = 2s, in-plane imaging resolution = $4 \times 4$ mm$^2$, in-plane field of view (FOV) = $256 \times 256$ mm$^2$, slice thickness = 5.5 mm with no gap, slice number = 23. The same rs-fMRI preprocessing procedures described in the Section 3.1 for adult brain was conducted on the second group of datasets.

Supp. Fig 2a shows the functional networks (FNs) generated by ICA, NNMF, Ncut, SCSC and RNcut. As indicated by red arrows, much fewer spurious patches were generated by the RNcut. Supp. Fig 2b show that statistically highest SI values were obtained with the RNcut. Supp. Fig 2c demonstrates that smaller number of isolated patches were generated by the RNcut compared to ICA, NNMF and Ncut. SCSC yielded even smaller number of isolated patches (Supp. Fig. 2c), but disrupted the shapes of underlying adult brain functional networks (Supp. Fig. 2a).

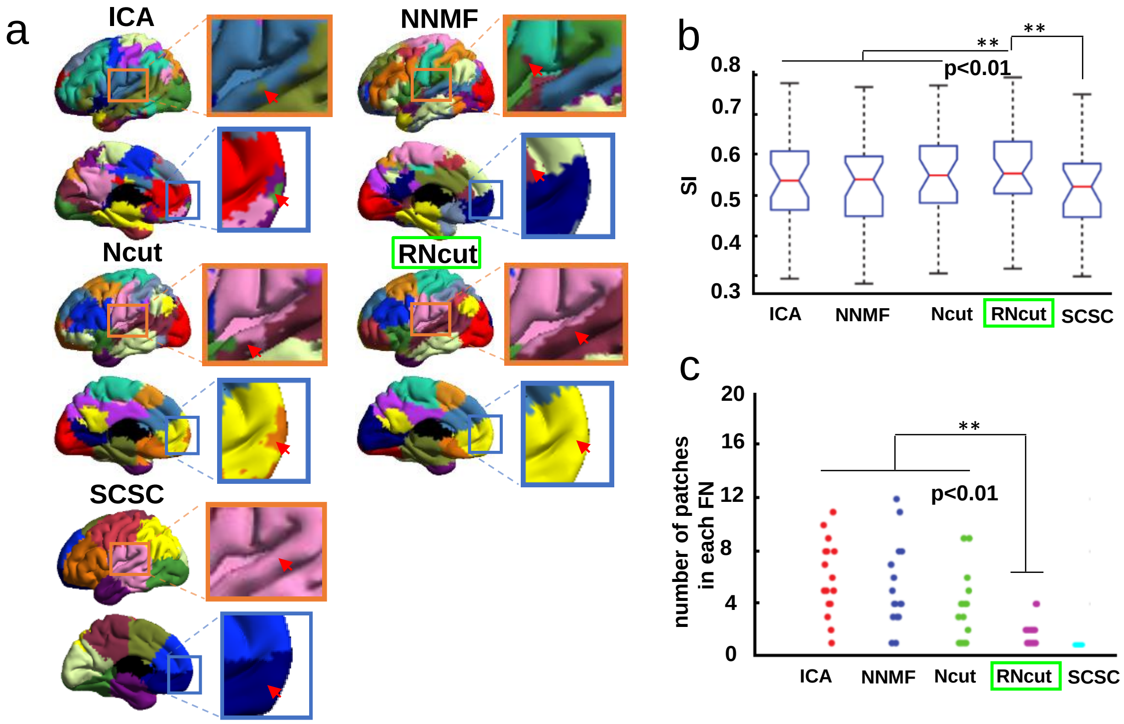


**Supplementary Figure 2: Parcellation of the functional networks of the adult group #2 brains using different methods. In panel a, the parcellation results of ICA, NNMF, Ncut, SCSC and RNcut demonstrate that fewer isolated spurious patches were obtained with RNcut, indicated by red arrows and highlighted with the enlarged boxes. Panel b shows significantly higher Silhouette Index (SI) value with RNcut method than all other competing methods. Panel c shows significantly smaller number of isolated patches in each functional network generated by RNcut method, compared to ICA, NNMF and Ncut. ** indicates p<0.01.**

<u>Intra-network connectivity</u>

Supp. Fig 3 shows intra-network connectivity in neonate brains based on the FNs parcellated by ICA, NNMF, Ncut, SCSC and RNcut. In Supp. Fig 3, high contrast aligned with the boundary of the primary sensorimotor region is clearly shown in the neonate brain intra-network connectivity map obtained with RNcut, but not shown in the maps obtained with other methods, especially ICA, NNMF or SCSC.

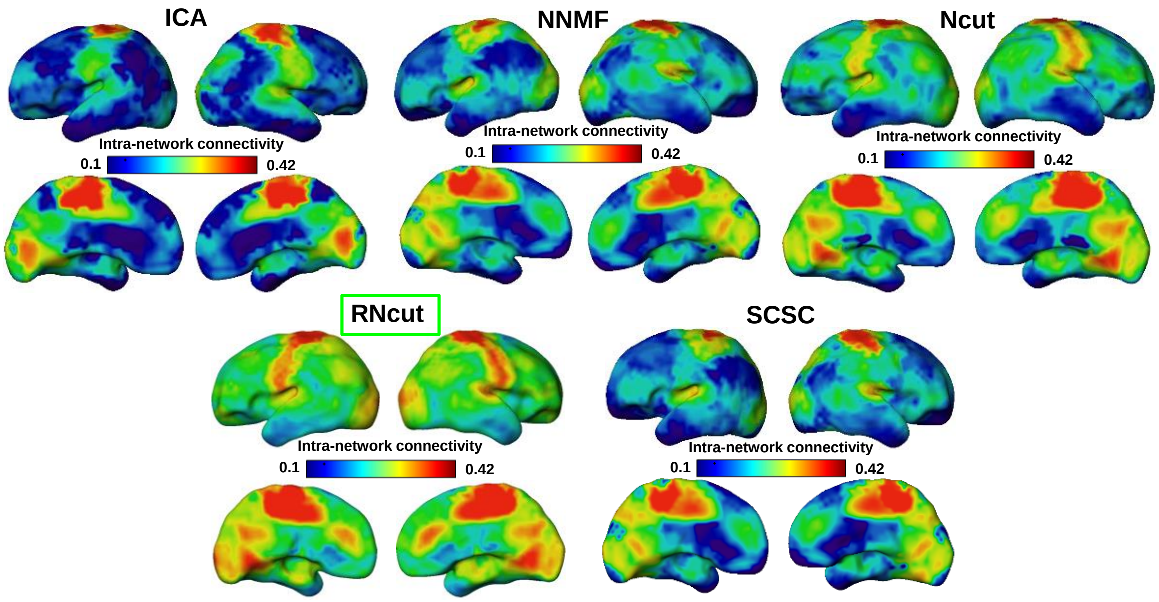


**Supplementary Figure 3: Intra-network connectivity of neonate brain using different methods. The intra-network connectivity map from RNcut reveals clear developmental pattern with high contrast aligned with the boundary of primary sensorimotor area.**
